# Olfactory chemosensation extends lifespan through TGF-β signaling and UPR activation

**DOI:** 10.1101/2022.10.12.511902

**Authors:** Evandro A. De-Souza, Maximillian A. Thompson, Rebecca C. Taylor

## Abstract

Animals rely on chemosensory cues to survive in pathogen-rich environments. In *C. elegans*, pathogenic bacteria are known to trigger aversive behaviors through neuronal perception, and to activate molecular defenses throughout the animal. This suggests that neurons may be able to coordinate the activation of organism-wide defensive responses upon pathogen perception. We find that exposure to volatile pathogen-associated compounds induces cell non-autonomous activation of the endoplasmic reticulum unfolded protein response (UPR^ER^) in peripheral tissues following *xbp-1* splicing in neurons. This odorant-induced UPR^ER^ activation is dependent upon transforming growth factor beta (TGF-β) signaling and leads to extended lifespan and enhanced clearance of toxic proteins. Our data suggest that the cell non-autonomous UPR^ER^ rewires organismal proteostasis in response to pathogen detection, pre-empting the arrival of proteotoxic stress. Thus, chemosensation of particular odors may be a novel way to manipulate stress responses and longevity.

## Main text

To adapt and survive, an organism must be able to detect and respond to environmental changes. In animals, this is mediated by the sensory nervous system, which activates defensive responses upon identification of hazards such as reduced oxygen availability, temperature increase or food shortage^1^. In addition, the detection of stress within cells can activate cellular stress responses such as the unfolded protein response of the endoplasmic reticulum (UPR^ER^), which respond to homeostatic imbalance by activating mechanisms that restore homeostasis^2^. As animals age, they lose this ability to recognize and respond to stress, resulting in increased mortality and age-related disease^1,3–5^. In particular, reduced activity of the IRE-1/XBP-1 signaling branch of the UPR^ER^ has been linked to brain aging and neurodegeneration, while genetic activation of XBP-1 can protect animals against proteotoxic insults^5,6^.

Recent evidence suggests that neurons can trigger the cell non-autonomous activation of cellular stress responses in peripheral tissues, leading to coordinated increases in organismal resilience and lifespan. Consistent with this, the genetic activation of the UPR^ER^ in a subset of neuronal or glial cells can extend lifespan in *C. elegans*, via neuronal signaling mechanisms that result in UPR^ER^ activation in distal tissues^7,8^. However, whether specific environmental situations or exogenous molecules can trigger the activation of the cell non-autonomous UPR^ER^ in wild-type animals remains unknown.

Olfactory perception of bacteria has been shown to alter gene expression in invertebrates^9^ and the immune response to *Pseudomonas spp* is associated with activation of the UPR^ER^ in *C. elegans*^3,10^ The smell of pathogenic bacteria can also sensitize the heat shock response in worms^11^, suggesting a possible link between olfaction and proteostasis. Therefore, we decided to investigate whether pathogen-associated odor could activate the cell non-autonomous UPR^ER^ in *C. elegans*. We exposed animals to a variety of odorant molecules secreted by pathogenic bacteria including *Pseudomonas aeruginosa* and *Staphylococcus aureus*^12^, and monitored the expression of *hsp-4p::GFP*, a transcriptional reporter of UPR^ER^ activation. Importantly, because the volatile molecules and the worms were placed on different plates, there was no direct contact between them (Fig. 1a). We observed that the UPR^ER^ could be activated in the intestine by exposure to three odorant molecules: 1-undecene, pyrrole, and 2-nonanone (Fig. 1b,c). Curiously, all three of these compounds had previously been linked to aversive behavioral responses in worms^13,14^ The chemical structures of 1-undecene and 2-nonanone are similar, both being hydrocarbons with a single carbonyl group. As 2-nonanone caused a degree of toxicity, we decided to focus upon 1-undecene in subsequent experiments.

**Fig. 1:**
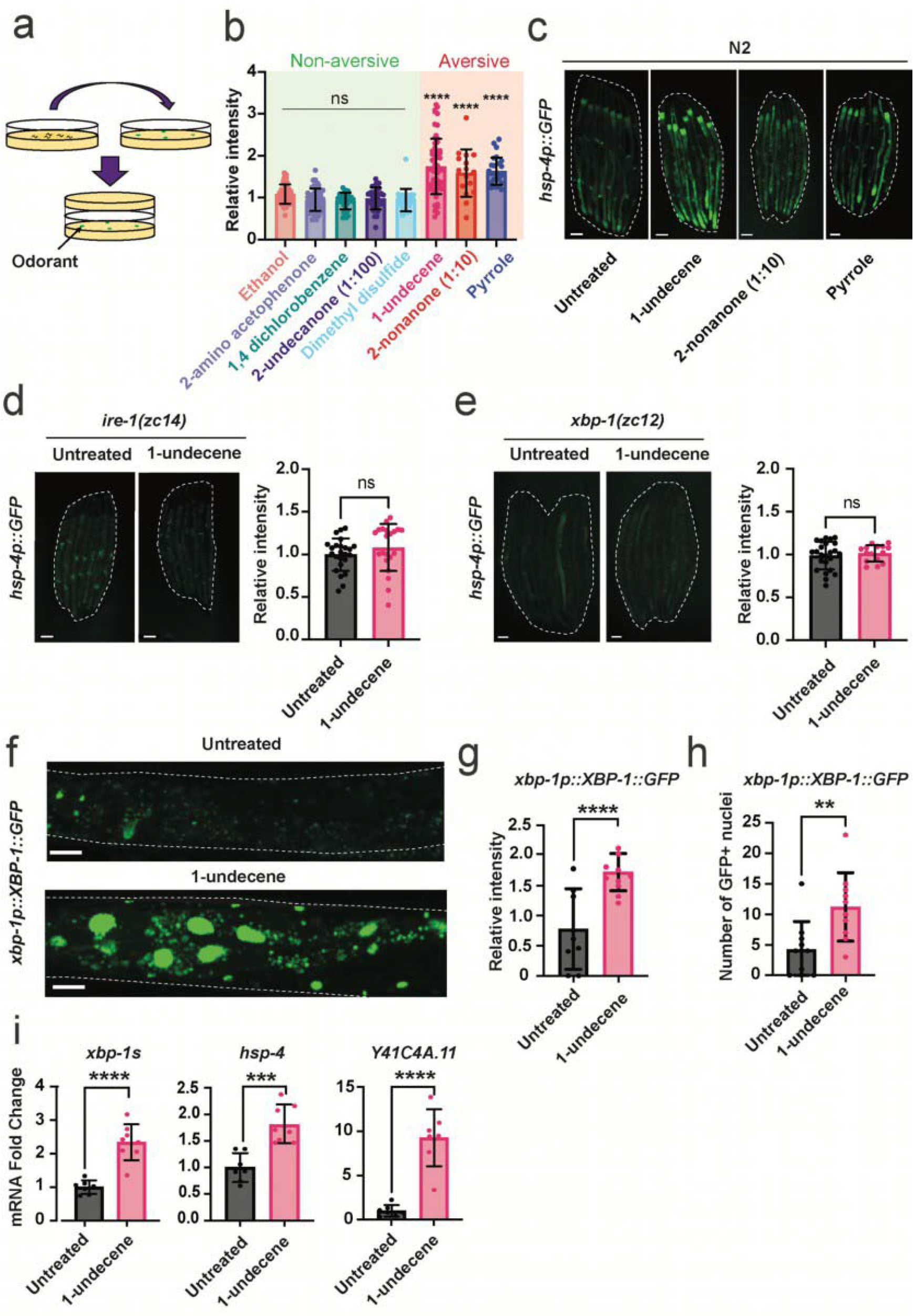
Pathogen-associated odor activates the IRE-1/XBP-1 branch of the UPR^ER^. **a,** Schematic showing the experimental setup for the odorant exposure assay. Briefly, around 30 young adult worms were sealed for 12 hours in NGM plates together with another NGM plate containing 4 spots of 3 μL of odorant. **b,** Quantification of *hsp-4p::GFP* expression was performed in ImageJ and data was normalized to untreated *hsp-4p::GFP* animals. This assay was independently performed three times with at least 10 worms per group. \*\*\*\**P*<0.0001 (one-way ANOVA with Dunnett’s multiple comparison test). **c,** Representative fluorescence microscopy images of worms untreated or exposed to 1-undecene, 2-nonanone (diluted 10x), and pyrrole for 12 hours. Scalebars, 200 μm. **d,e,** Representative fluorescence microscopy images and quantification of *hsp-4p::GFP* fluorescence in **d,** *ire-1(zc14*) and **e**, *xbp-1(zc12*) worms with or without exposure to 1-undecene odor for 12 hours. ns, not significant (unpaired Student’s t test). These experiments were repeated at least three times with more than 10 worms per group. Scalebars, 200 μm. **f,** Representative image, **g,** quantification of fluorescence, and **h,** number of GFP-positive nuclei in the intestine of worms expressing an *xbp-1p::xbp-1::GFP* transgene with or without exposure to 1-undecene for 8 hours. Scalebars, 200 μm. This experiment was repeated three times with 10 worms per group. \*\*\*\**P*<0.0001, \*\**P*<0.01 (unpaired Student’s t test). **i,** mRNA levels of *xbp-1s, hsp-4*, and *Y41C4A.11* were measured by qRT-PCR in animals exposed to 1-undecene for 8 hours relative to untreated worms. n=7-8 biological replicates. Ns, not significant, \*\*\*\**P*<0.0001, \*\*\**P*<0.001 (unpaired Student’s t test).

We found that mutation of the UPR^ER^ regulators *ire-1* or *xbp-1* abolished UPR^ER^ activation by 1-undecene odor, indicating that the IRE-1/XBP-1 signaling pathway is essential for activation of the UPR^ER^ by this compound (Fig. 1d,e). Consistent with this, an XBP-1s::GFP splicing reporter that expresses XBP-1s::GFP from an *xbp-1p::xbp-1::GFP* transgene only when *xbp-1* mRNA is spliced by IRE-1^8^ revealed an increase in XBP-1s::GFP fluorescence within the intestinal cells of animals exposed to 1-undecene (Fig. 1f,g). Furthermore, we observed a significant increase in transcript levels of spliced *xbp-1* and two XBP-1 target genes (*hsp-4* and *Y41C4A.11*), confirming activation of the IRE-1/XBP-1 pathway by 1-undecene (Fig. 1h). Interestingly, we were unable to detect activation of other cellular stress response pathways, including nuclear DAF-16 localization and *hsp-16.2* (heat shock response) or *hsp-6* (mitochondrial UPR) upregulation, suggesting that the UPR^ER^ is specifically activated by pathogen-associated odor (Extended Data Fig. 1). Finally, a recent study found that the *C. elegans* immune system can also be activated by olfactory perception of 1-undecene^15^. However, odor-induced UPR^ER^ activation is unlikely to be a downstream consequence of immune response activation, as animals with a mutation in the key immunity transcription factor *zip-2* still showed UPR^ER^ activation in response to 1-undecene (Extended Data Fig. 2a).

Previous work from our group and others has demonstrated that neuronal signaling can activate the UPR^ER^ in peripheral tissues such as the intestine^3,16^. We wondered whether signals produced by the nervous system were also responsible for odor-induced UPR^ER^ activation. We observed that animals exposed to pathogen-associated odor had a significant increase in both the number and fluorescence intensity of XBP-1s::GFP-positive neuronal cells (Fig. 2a and Extended Data Fig. S2b). To establish whether UPR^ER^ activation arising from 1-undecene exposure was cell non-autonomous in nature, we tested the dependency of this effect on the neuronal signaling regulators *unc-31* and *unc-13* – mutations in the former blocking release of neuropeptides from dense core vesicles, and in the latter preventing the release of a range of signaling molecules including small-molecule neurotransmitters^3,7^. We observed that the *hsp-4p::GFP* reporter was activated in the intestine of *unc-31(e928*) mutant animals, suggesting that olfactory cues do not require neuropeptide signaling to activate the UPR^ER^ (Fig. 2b). In contrast, the *unc-13(e450*) mutation entirely inhibited activation of the UPR^ER^ in the periphery, demonstrating that a non-neuropeptide neuronal signal is involved in cell non-autonomous UPR^ER^ activation by exposure to 1-undecene (Fig. 2c). Importantly, mutation of *unc-13* does not prevent animals from responding to cell-autonomous ER stress, as *hsp-4p::GFP* is still activated in animals exposed to RNAi against *pdi-2* (Extended Data Fig. S2c)^17^.

**Fig. 2:**
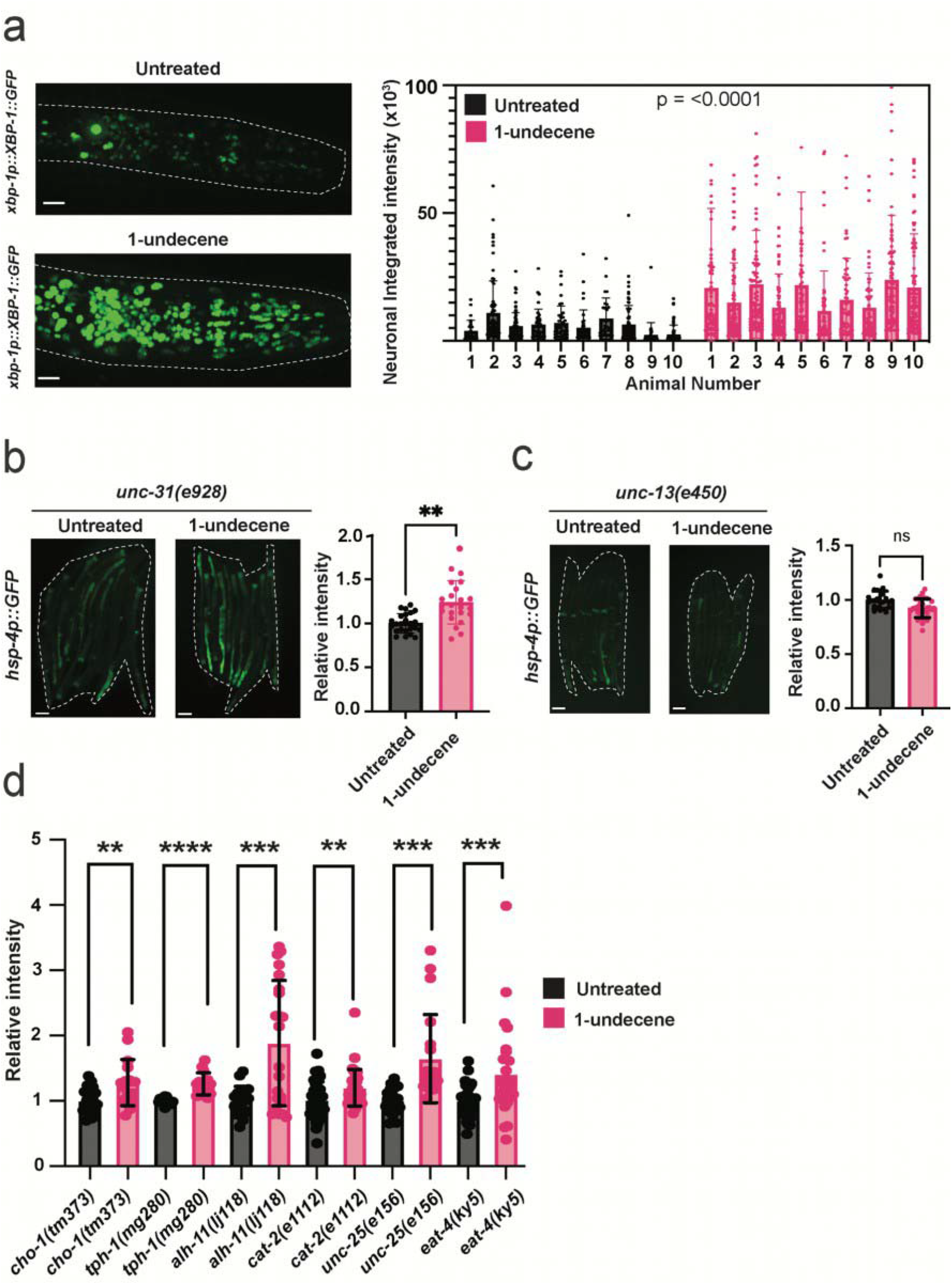
Neuronal signaling is required for downstream UPR^ER^ activation by 1-undecene exposure. **a,** Representative image and quantification of GFP-positive neuronal cells in worms expressing an *xbp-1p::xbp-1::GFP* transgene with or without exposure to 1-undecene for 8 hours. This experiment was repeated three times with 10 worms per group. (unpaired Student’s t test). Scalebars, 10 μm. **b, c,** Representative fluorescence microscopy images and quantification of *hsp-4p::GFP* fluorescence in **b,** *unc-31(e928*) and **c,** *unc-13(e450*) with or without exposure to 1-undecene odor for 12 hours. ns, not significant, \*\**P*<0.01 (unpaired Student’s t test). These experiments were repeated at least three times with more than 10 worms per group. Scalebars, 200 μm. **d,** Quantification of *hsp-4p::GFP* fluorescence in *cho-1(tm373), tph-1(mg280), alh-1(ij118), cat-2(e1112), unc-25(e156), eat-4(ky5*). Intensity was normalized to untreated animals for each mutant strain. \*\**P*<0.01, \*\*\**P*<0.01, \*\*\*\**P*<0.001 (unpaired Student’s t test comparison between untreated and 1-undecene).

The Gα protein ODR-3 was previously shown to be required for activation of the immune response by 1-undecene^15^. We therefore asked whether ODR-3 is also required for 1-undecene-induced UPR^ER^ activation. However, we observed a full *hsp-4p::GFP* response in an *odr-3* null background, suggesting that this gene is not required for UPR^ER^ activation (Extended Data Fig. S3a). In addition, tyramine synthesis is necessary for cell non-autonomous UPR^ER^ activation in strains constitutively expressing neuronal *xbp-1s*^8^. Unexpectedly, however, we found that *tdc-1*, a gene essential to synthesize tyramine, was not required for activation of *hsp-4p::GFP* in strains exposed to 1-undecene (Extended Data Fig. S3b). We also ruled out the possibility that the CEPsh glia, another cell type implicated in cell non-autonomous UPR^ER^ activation^7^, were involved in this response, as animals in which these cells were genetically ablated still displayed an increase in *hsp-4p::GFP* levels following 1-undecene exposure (Extended Data Fig. S3c). We then tested mutants that fail to synthesize a variety of neurotransmitters, including dopamine, serotonin, GABA, glutamate, choline, and betaine, for their ability to activate the UPR^ER^ in response to 1-undecene exposure, but were unable to identify a role for any of these molecules (Fig. 2d and Extended Data Fig. S4).

Worms avoid food containing pathogenic bacteria through aversive olfactory learning^18^. The same form of aversive behavior is seen in animals exposed to pathogen-associated molecules^19,20^. One signaling molecule known to play a key role in the neuronal circuits that govern these behaviors is transforming growth factor-beta (TGF-β)^19,21^. DAF-7, a worm homologue of TGF-β, is also necessary for the avoidance of 2-nonanone^22^, a molecule whose odor induced UPR^ER^ activation in our initial odorant screen (Fig. 1b). We therefore asked whether DAF-7/TGF-β is required for UPR^ER^ activation by 1-undecene. Strikingly, we found that *daf-7* was indeed necessary for UPR^ER^ activation following 1-undecene exposure (Fig. 3a). In addition, a mutation in a specific DAF-7 receptor, *daf-1(m40*), also completely inhibited odorant-induced UPR^ER^ activation (Fig. 3b). Importantly, DAF-1 is expressed in the RIM/RIC interneurons, and our previous work has shown that UPR^ER^ activation in these neurons is sufficient to drive inter-tissue intestinal UPR^ER^ activation^19,23^. DAF-7 is primarily expressed in the ASI chemosensory neurons, and animals exposed to *P. aeruginosa* exhibit increased expression of *daf-7*^19^. We therefore asked whether *daf-7* expression could also be elevated by chemosensation of 1-undecene. Indeed, *daf-7* mRNA levels were upregulated upon 1-undecene exposure (Fig. 3c). To confirm this, we also employed a *daf-7p::Venus* fluorescent reporter transgene, and observed an increase in the expression of *daf-7* in the ASI neurons upon treatment with 1-undecene (Fig. 3d).

**Fig. 3:**
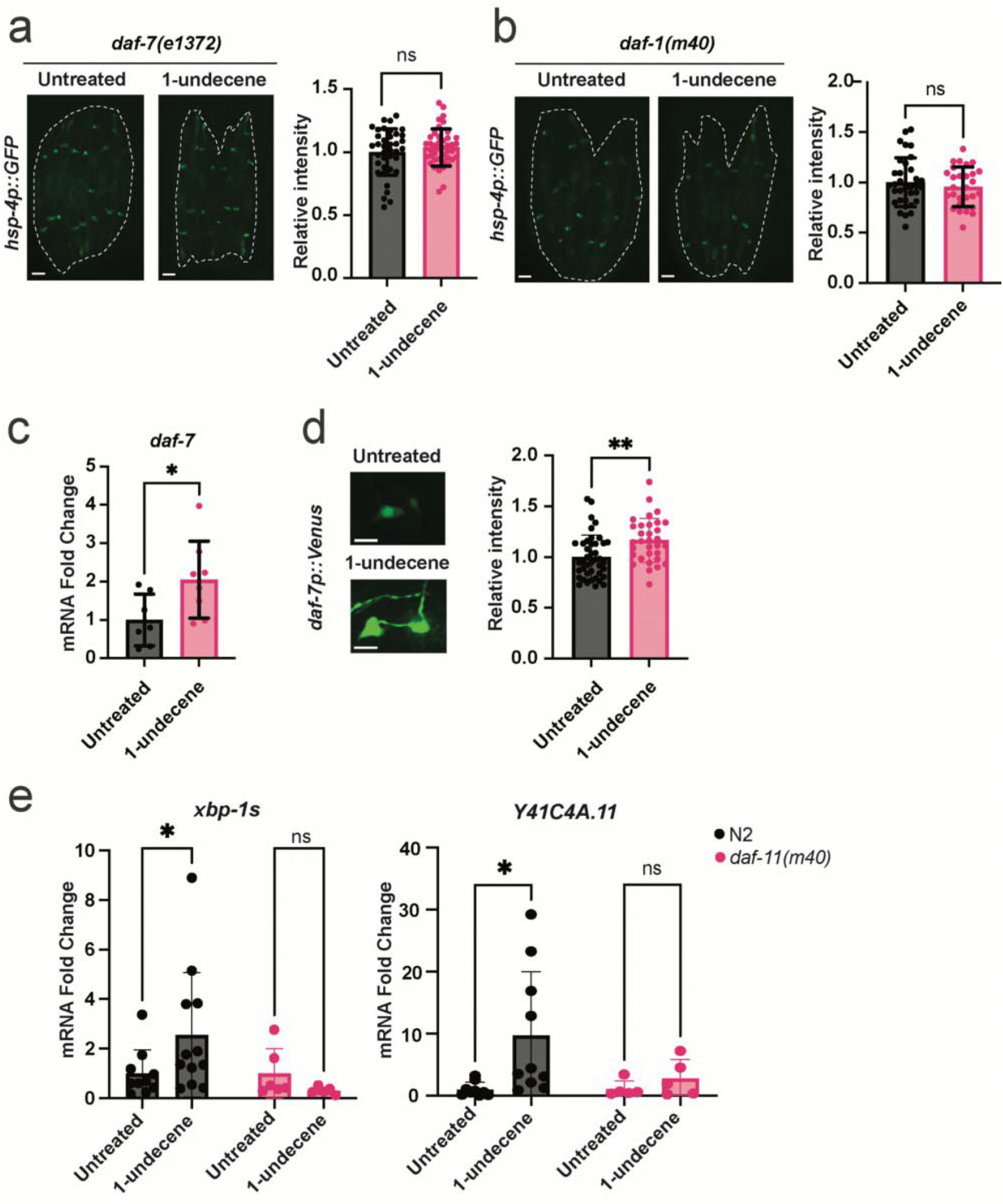
DAF-7/TGF-β signaling is required for odor-induced UPR^ER^ activation. **a, b,** Representative fluorescence microscopy images and quantification of *hsp-4p::GFP* fluorescence in **a,** *daf-7(e1372*) and **b,** *daf-1(m40*) strains with or without exposure to 1-undecene for 12 hours. Each experiment was repeated more than three times with at least 10 worms per group. Ns, not significant (unpaired Student’s t test). Scalebars, 200 μm**. c,** mRNA levels of *daf-1* were measured by qRT-PCR in animals exposed to 1-undecene for 8 hours relative to untreated worms. n=7-8 biological replicates. \**P*<0.05 (unpaired Student’s t test). **d,** Representative fluorescence microscopy images and quantification of *daf-7p::Venus* fluorescence in ASI neurons after worms were exposed or not exposed to 1-undecene odor for 12 hours. This experiment was repeated three times with at least 10 worms per group. \*\**P*<0.01 (unpaired Student’s t test). Scalebars, 7 μm. **e,** mRNA levels of *xbp-1s* and *Y41C4A.11* were measured by qRT-PCR in animals exposed to 1-undecene for 8 hours relative to untreated worms. n=5-12 biological replicates. Ns, not significant, \**P*<0.05 (Two-Way ANOVA with Tukey’s multiple comparison test).

Expression levels of *daf-7* have been previously linked to activation of the guanylate cyclase DAF-11 in ASI neurons during starvation^24^. We therefore asked whether DAF-11 is also required for UPR^ER^ activation upon 1-undecene exposure, and observed that DAF-11 was indeed necessary for transcriptional upregulation of *xbp-1s* and the XBP-1 target gene *Y41C4A.11* (Fig. 3e). This therefore suggests that DAF-11 is involved in the neuronal perception of 1-undecene odor and subsequent UPR^ER^ activation. Thus, our data implicates a TGF-β signaling circuit in connecting the recognition of pathogen-related odorants to inter-tissue regulation of the UPR^ER^.

Activation of cellular stress responses has been associated with increased lifespan and improved resistance to disease-associated toxic protein species^6,25,26^. This prompted us to ask whether 1-undecene exposure on the first day of adulthood could impact organismal lifespan and proteostasis. Excitingly, 1-undecene-exposed animals consistently had significantly longer lifespans than untreated animals (Fig. 4a and Supplementary Table 1). This increase in survival was dependent on *xbp-1* (Fig. 4b and Supplementary Table 1), suggesting that 1-undecene odor extends lifespan through the activation of the UPR^ER^. To examine the impact of pathogen-related odor on a *C. elegans* model of neurodegeneration-associated proteotoxicity, we measured levels of YFP-tagged polyglutamine (polyQ) repeats in different tissues of the animal following 1-undecene exposure at day 1 of adulthood. Remarkably, 1-undecene induced a consistent decrease in levels of polyQ in all tissues examined (intestine, muscle, and neurons), suggesting that 1-undecene-induced UPR^ER^ activation enhances clearance of toxic proteins across the animal (Fig. 4c). These results therefore suggest a model in which the neuronal perception of an odorant molecule can influence organismal proteostasis and lifespan through TGF-β signaling and UPR^ER^ activation (Fig. 4d).

**Fig. 4:**
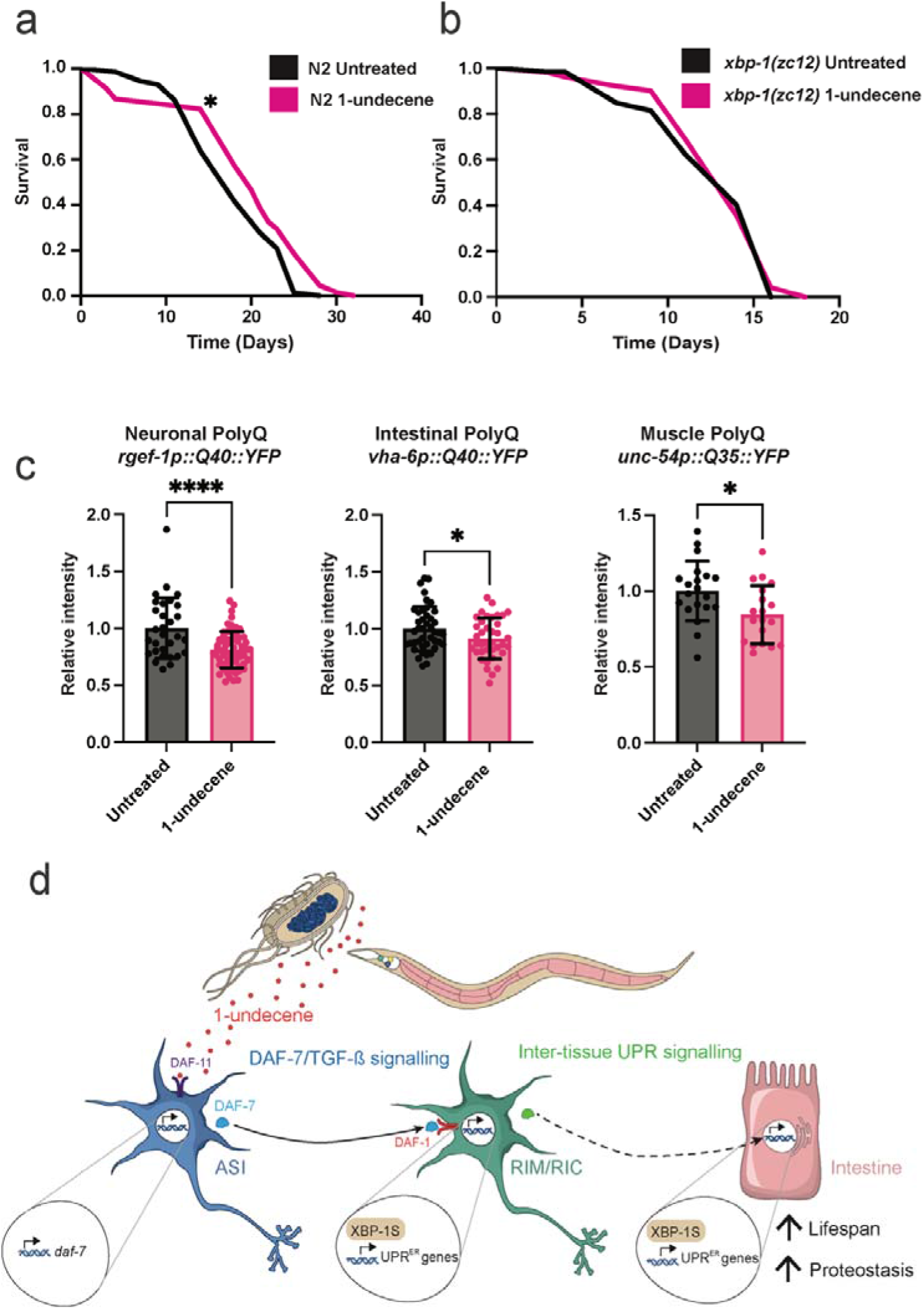
1-undecene odor increases *C. elegans* lifespan and reduces polyglutamine accumulation. **a,b,** Lifespan analysis of **a,** N2 wild-type and **b,** *xbp-1(zc12*) animals with or without exposure to 1-undecene for 24 hours at day 1 of adulthood. Graphs are plotted as Kaplan-Meier survival curves. N=100-120 animals in each group. \**P*<0.05 (Mantel-Cox log-rank test). **c,** Animals expressing polyglutamine:YFP repeats in neurons, intestine or body wall muscle were exposed to 1-undecene for 12 hours at day 1 of adulthood and imaged 72 hours after treatment. YFP levels were quantified using ImageJ and normalized to untreated animals. \*\*\*\**P*<0.0001, \**P*<0.05 (unpaired Student’s t test)**. d,** Model showing that olfactory stimulation, through exposure to 1-undecene, can activate the cell non-autonomous UPR^ER^ in *C. elegans* via TGF-β signaling.

Previous studies have reported the cell non-autonomous activation of the UPR^ER^ by signals from neurons and glia. In each case, however, transgenes driving *xbp-1s* have been used to achieve this activation and the evolutionary logic for the development of such systems has been unclear. Here we demonstrate that *C. elegans* are capable of triggering a cell non-autonomous UPR^ER^ without such transgenes, in response to several odorant molecules that trigger an aversive behavioral response and are secreted by pathogenic bacteria. We reason that the cell non-autonomous UPR^ER^ may therefore have evolved in order to enable the animal to enhance defensive mechanisms in anticipation of the increased translation associated with an immune response, or the direct proteostatic challenge of the pathogen itself. Animals that constitutively activate a PMK-1-driven immune response require *xbp-1* to survive the demands imposed by an active immune system^**10**^, suggesting that the requirement for enhanced UPR^ER^ capacity is of critical importance in conditions that require an immune response.

While the action of 1-undecene on *C. elegans* is likely a specific interaction informed by the complex evolutionary relationship between pathogen and host, there is existing evidence to support the idea that the broader principle underlying this type of cell non-autonomous UPR^ER^ activation may be conserved. The most striking is the finding that in mice, the sensory perception of food activates POMC (Proopiomelanocortin expressing) neurons, resulting in hepatic *xbp-1* splicing as a predictive physiological response in anticipation of food consumption^27^. It has also been demonstrated that driving *xbp-1s* genetically in murine POMC neurons is sufficient to increase hepatic *xbp-1s* levels via a cell non-autonomous mechanism^16^. There are significant similarities between the roles of ASI neurons in the worm and the hypothalamus and POMC neurons in mice, and ASI neurons have been referred to as the “putative hypothalamus” of the worm due to these similar functions^28^. ASI neurons regulate food intake and food seeking behaviour through the action of DAF-7/TGF-β^29^. Similarly, POMC is expressed in subsets of cells including neurons in the arcuate nucleus of the hypothalamus^30^, and POMC neurons also regulate food intake and energy expenditure via locomotion in some contexts^31^. Furthermore, expression of the TGF-β antagonist Smad7 in POMC neurons regulates peripheral glucose metabolism, suggesting that TGF-β signalling is also important for POMC neurons to achieve anticipatory, cell non-autonomous effects in the periphery^32^. These mammalian studies suggest that major interactions in the pathway we describe here are likely to be conserved in mammalian systems.

Although earlier studies have shown that food-associated odor can prevent lifespan extension induced by caloric restriction^**33,34**^, we believe this study is the first demonstration that the perception of a specific odorant molecule can increase the lifespan of an animal. It has been noted recently^**35**^ that a great many mechanisms which regulate aging in model organisms include cell non-autonomous protective pathways that are subject to neuronal control, often by sensory neurons. Dietary restriction-mediated longevity requires the UPR^ER^^**36,37**^ as well as functional ASI neurons expressing *daf-7* **^38,39^**, and is regulated by olfactory perception^**40**^. Furthermore, cell non-autonomous regulation of not just the UPR^ER^, but also the mitochondrial UPR^**41**^, heat shock response^**42**^, AMP-activated protein kinase (AMPK)^**43**^, and target of rapamycin complex 1 (TORC1)^**44**^, as well as lifespan regulation by temperature^**45**^ and the hypoxia response^**46**^ are all similarly orchestrated, with signals originating in sensory neurons leading via cell non-autonomous routes to regulation of pro-longevity pathways. Here, we show that direct activation of chemosensory neurons by the ligands they sense can extend lifespan. We therefore speculate that directly manipulating the activity of sensory neurons via their sensory inputs and/or corresponding receptors may be a novel way to activate these prolongevity pathways.

Finally, mounting evidence suggests that Ire1/Xbp1 activity is highly correlated with the pathophysiology observed in various neurodegenerative disorders in animal models, including Alzheimer’s, Parkinson’s and Huntington’s diseases, and age-associated decline in the activation of this pathway may be associated with disease progression^**47-49**^. Activation of the UPR^ER^ through stimulation of sensory pathways by olfactory compounds may therefore represent a promising strategy to prevent the disease-related proteostasis collapse associated with aging.

## Acknowledgments

We are grateful to the MRC LMB Visual Aids department for assistance with figures. Some *C. elegans* strains were provided by Prof. Andrew Dillin (UC Berkeley), Dr. William Schafer (MRC-LMB), Dr. Jennifer Tullet (University of Kent), and the Caenorhabditis Genetics Center (CGC), which is funded by the NIH Office of Research Infrastructure Programs (P40 OD010440). This work was supported by the Medical Research Council (R.C.T.) and by the European Union’s Horizon 2020 research and innovation programme under the Marie Skłodowska-Curie grant agreement number 894039 (E.A.D.).

## Author contributions

Conceptualization: EAD, MAT, RCT

Methodology: EAD, MAT, RCT

Investigation: EAD, MAT, RCT

Visualization: EAD, MAT, RCT

Funding acquisition: EAD, RCT

Project administration: RCT

Supervision: RCT

Writing – original draft, review & editing: EAD, MAT, RCT

## Methods

### *C. elegans* strains and maintenance

Strains were made in the course of this study, provided by the CGC, or kindly gifted by other labs. A list of strains used in this work can be found in Supplementary Table S2. We used the CGC Bristol N2 hermaphrodite stock as our wild-type. Worms were maintained at 20°C on NGM (nematode growth medium) plates seeded with *Escherichia coli* OP50 using standard techniques^50^ For RNAi by feeding^51^, NGM plates were supplemented with 1 mM IPTG and 100 μg/mL carbenicillin and then seeded with HT115 bacteria harboring L4440 empty vector or the RNAi of interest. All RNAi used are from the Ahringer RNAi library (Source Bioscience) and were confirmed by sequencing.

### Transgenic strain construction

The *odr-3(rms31*) mutant was generated by CRISPR using a dual crRNA *dpy-10* co-crispr strategy and a custom protocol based on previous methods^52,53^ and optimization for our lab. Briefly, 1 μL of 320 μM solution of each crRNA and 0.5 μL of *dpy-10* crRNA (50 μM) was annealed to 0.4 μL of 100 μM tracrRNA (IDT) by heating to 95 °C in a PCR machine and cooling to 4 °C at 0.1 °C/sec. 0.5 μL of Cas9 protein (Invitrogen) was then added and the mixture was incubated for 10min at 37 °C. 0.5 μL of 100 μM stock of each repair template (target and *dpy-10*) and the solution made up to 20 μL with DPEC water. This mix was centrifuged for 30 min at 13,000 rpm at 4 °C before injection. Oligonucleotides used in this paper can be found in Supplementary Table S3 and S4.

### Epifluorescence microscopy

To investigate the effect of 1-undecene on reporter transgene expression (*e.g. hsp-4p::gfp*), worms were exposed to 1-undecene odor for 12 hours in plates sealed with Parafilm M^®^, and then immobilized with 20 mM of sodium azide (Sigma) and imaged using a Leica M205 FA microscope. To image worms expressing polyQ::YFP, worms were exposed to 1-undecene for 8 hours on day 1 of adulthood and imaged on day 4 of adulthood. For DAF-16::GFP analysis, worms were scored based on the subcellular localization of GFP in intestinal cells as described before^24^. Worms were randomly selected from a synchronized population before imaging. Fluorescence values (mean intensity) were obtained by analyzing microscope images on ImageJ or Fiji.

### Confocal microscopy

Worms were immobilized with 20 mM of sodium azide (Sigma) and mounted on a 2% agarose pad. Animals were imaged on an LSM 710 confocal microscope using the 40x and 63x oil immersion objectives and on an Andor Revolution spinning disk microscope using the 20x and 60x water immersion objectives. All images were analyzed using ImageJ or Fiji.

### RNA extraction and qRT-PCR

Approximately 300 young adult animals were collected with M9 after being exposed or not to 1-undecene for 8 hours. Trizol was added to samples, which were immediately frozen in liquid nitrogen. RNA isolation was carried out using the Directzol RNA Miniprep kit (Zymo Research) following the manufacturer’s instructions. RNA was quantified by Nanodrop. 1 μg of RNA was used for cDNA synthesis with the QuantiTect Reverse Transcription kit (Qiagen). Samples were diluted 2.5x after cDNA synthesis and SYBR™ Select Master Mix (Applied Biosystems) was used for quantitative RT-PCR on a Vii7 Real-Time PCR machine (ThermoFisher Scientific) to quantify alterations in the transcript level of genes of interest. Data were analyzed using the comparative 2^ΔΔ^Ct method. A list of primers used in this work can be found in Table S3.

### Survival assays

Approximately 100 worms were exposed or not to 1-undecene odor for 24 hours. Worms were then placed on NGM plates containing 100 μg/mL FUDR and seeded with *E. coli* OP50, and were kept at 20°C. Animals were monitored as alive or dead every second day by a blinded investigator and data was analyzed on GraphPad Prism 8 software.

### Statistics

Statistical analysis was performed using GraphPad Prism 8 software. All bar graphs show the mean with error bars representing standard deviation. Appropriate tests for each experiment were chosen and are described (including tests for multiple comparisons etc) in the figure legends. Where used, “n” is immediately defined. Information regarding the number of repeats, number of animals per repeat and the results of the statistical tests performed are given in the figure legends.

### Data availability

All data reported in this paper will be shared by the lead contact upon request. This paper does not report original code. Any additional information required to reanalyze the data reported in this paper is available from the lead contact upon request.

## References

1. Linford, N. J., Kuo, T.-H., Chan, T. P. & Pletcher, S. D. Sensory perception and aging in model systems: from the outside in. Annu. Rev. Cell Dev. Biol. 27, 759–85 (2011).

2. Walter, P. & Ron, D. The unfolded protein response: from stress pathway to homeostatic regulation. Science 334, 1081–6 (2011).

3. Taylor, R. C. & Dillin, A. XBP-1 Is a Cell-Nonautonomous Regulator of Stress Resistance and Longevity. Cell 153, 1435–1447 (2013).

4. Labbadia, J. & Morimoto, R. I. Repression of the Heat Shock Response Is a Programmed Event at the Onset of Reproduction. Mol. Cell 59, 639–650 (2015).

5. Cabral-Miranda, F. et al. Control of mammalian brain aging by the unfolded protein response (UPR). bioRxiv (2020) doi:10.1101/2020.04.13.039172.

6. Imanikia, S., Özbey, N. P., Krueger, C., Casanueva, M. O. & Taylor, R. C. Neuronal XBP-1 Activates Intestinal Lysosomes to Improve Proteostasis in C. elegans. Curr. Biol. CB 29, 2322–2338.e7 (2019).

7. Frakes, A. E. et al. Four glial cells regulate ER stress resistance and longevity via neuropeptide signaling in C. elegans. Science 367, 436–440 (2020).

8. Özbey, N. P. et al. Tyramine Acts Downstream of Neuronal XBP-1 s to Coordinate Inter-tissue UPRER Activation and Behavior in C. elegans. Dev. Cell 55, 754–770.e6 (2020).

9. Masuzzo, A., Montanari, M., Kurz, L. & Royet, J. How Bacteria Impact Host Nervous System and Behaviors: Lessons from Flies and Worms. Trends Neurosci. 43, 998–1010 (2020).

10. Richardson, C. E., Kooistra, T. & Kim, D. H. An essential role for XBP-1 in host protection against immune activation in C. elegans. Nature 463, 1092–1095 (2010).

11. Ooi, F. K. & Prahlad, V. Olfactory experience primes the heat shock transcription factor HSF-1 to enhance the expression of molecular chaperones in C. elegans. Sci. Signal. 10, (2017).

12. Filipiak, W. et al. Molecular analysis of volatile metabolites released specifically by Staphylococcus aureus and Pseudomonas aeruginosa. BMC Microbiol. 12, 113 (2012).

13. L’Etoile, N. D. & Bargmann, C. I. Olfaction and odor discrimination are mediated by the C. elegans guanylyl cyclase ODR-1. Neuron 25, 575–86 (2000).

14. Prakash, D., Siddiqui, R., Chalasani, S. H. & Singh, V. Pyrrole produced by Pseudomonas aeruginosa influences olfactory food choice of Caenorhabditis elegans. bioRxiv 1–18 (2022).

15. Prakash, D. et al. 1-Undecene from Pseudomonas aeruginosa is an olfactory signal for flight-or-fight response in Caenorhabditis elegans. EMBO J. 40, (2021).

16. Williams, K. W. et al. Xbp1s in Pomc Neurons Connects ER Stress with Energy Balance and Glucose Homeostasis. Cell Metab. 20, 471–482 (2014).

17. Eletto, D., Eletto, D., Dersh, D., Gidalevitz, T. & Argon, Y. Protein disulfide isomerase A6 controls the decay of I REW1α signaling via disulfide-dependent association. Mol. Cell 53, 562–576 (2014).

18. Zhang, Y., Lu, H. & Bargmann, C. I. Pathogenic bacteria induce aversive olfactory learning in Caenorhabditis elegans. Nature 438, 179–84 (2005).

19. Meisel, J. D., Panda, O., Mahanti, P., Schroeder, F. C. & Kim, D. H. Chemosensation of bacterial secondary metabolites modulates neuroendocrine signaling and behavior of C. elegans. Cell 159, 267–80 (2014).

20. Beale, E., Li, G., Tan, M.-W. & Rumbaugh, K. P. Caenorhabditis elegans senses bacterial autoinducers. Appl. Environ. Microbiol. 72, 5135–7 (2006).

21. Moore, R. S., Kaletsky, R. & Murphy, C. T. Piwi/PRG-1 Argonaute and TGF-β Mediate Transgenerational Learned Pathogenic Avoidance. Cell 177, 1827–1841.e12 (2019).

22. Harris, G. et al. Molecular and cellular modulators for multisensory integration in C. elegans. PLOS Genet. 15, e1007706 (2019).

23. Özbey, N. P. et al. Tyramine Acts Downstream of Neuronal XBP-1s to Coordinate Inter-tissue UPRER Activation and Behavior in C. elegans. Dev. Cell 55, 754–770.e6 (2020).

24. Tataridas-Pallas, N. et al. Neuronal SKN-1B modulates nutritional signalling pathways and mitochondrial networks to control satiety. PLOS Genet. 17, e1009358 (2021).

25. Balch, W. E., Morimoto, R. I., Dillin, A. & Kelly, J. W. Adapting proteostasis for disease intervention. Science 319, 916–9 (2008).

26. Klaips, C. L., Jayaraj, G. G. & Hartl, F. U. Pathways of cellular proteostasis in aging and disease. J. Cell Biol. 217, 51–63 (2018).

27. Brandt, C. et al. Food Perception Primes Hepatic ER Homeostasis via Melanocortin-Dependent Control of mTOR Activation. Cell 175, 1321–1335.e20 (2018).

28. Tullet, J. M. a et al. Direct inhibition of the longevity promoting factor SKN-1 by insulin-like signaling in C. elegans. Cell 132, 1025–1038 (2008).

29. Gallagher, T., Kim, J., Oldenbroek, M., Kerr, R. & You, Y.-J. ASI Regulates Satiety Quiescence in C. elegans. J. Neurosci. 33, 9716–9724 (2013).

30. Yang, Y., Atasoy, D., Su, H. H. & Sternson, S. M. Hunger states switch a flipflop memory circuit via a synaptic AMPK-dependent positive feedback loop. Cell 146, 992–1003 (2011).

31. Zhan, C. et al. Acute and Long-Term Suppression of Feeding Behavior by POMC Neurons in the Brainstem and Hypothalamus, Respectively. J. Neurosci. 33, 3624–3632 (2013).

32. Yuan, F. et al. Overexpression of Smad7 in hypothalamic POMC neurons disrupts glucose balance by attenuating central insulin signaling. Mol. Metab. 42, 101084 (2020).

33. Libert, S. et al. Regulation of Drosophila life span by olfaction and food-derived odors. Science 315, 1133–7 (2007).

34. Park, S. et al. Diacetyl odor shortens longevity conferred by food deprivation in C. elegans via downregulation of DAF-16/FOXO. Aging Cell 20, e13300 (2021).

35. Miller, H. A., Dean, E. S., Pletcher, S. D. & Leiser, S. F. Cell non-autonomous regulation of health and longevity. eLife 9, e62659 (2020).

36. Li, P. W., Karpac, J., Jasper, H. & Kapahi, P. Intestinal IRE1 is required for increased triglyceride metabolism and longer lifespan under Dietary Restriction. 17, 1207–1216 (2016).

37. Matai, L. et al. Dietary restriction improves proteostasis and increases life span through endoplasmic reticulum hormesis. Proc. Natl. Acad. Sci. U. S. A. 116, 17383–17392 (2019).

38. Bishop, N. a & Guarente, L. Two neurons mediate diet-restriction-induced longevity in C. elegans. Nature 447, 545–549 (2007).

39. Fletcher, M. & Kim, D. H. Age-Dependent Neuroendocrine Signaling from Sensory Neurons Modulates the Effect of Dietary Restriction on Longevity of Caenorhabditis elegans. 1–15 (2017) doi:10.1371/journal.pgen.1006544.

40. Zhang, B., Jun, H., Wu, J., Liu, J. & Xu, X. Z. S. Olfactory perception of food abundance regulates dietary restriction-mediated longevity via a brain-to-gut signal. Nat. Aging 1, 255–268 (2021).

41. Durieux, J., Wolff, S. & Dillin, A. The Cell-Non-Autonomous Nature of Electron Transport Chain-Mediated Longevity. Cell 144, 79–91 (2011).

42. Prahlad, V., Cornelius, T. & Morimoto, R. I. Regulation of the Cellular Heat Shock Response in Caenorhabditis elegans by Thermosensory Neurons. Science 320, 811–814 (2008).

43. Burkewitz, K. et al. Neuronal CRTC-1 Governs Systemic Mitochondrial Metabolism and Lifespan via a Catecholamine Signal. Cell 160, 842–855 (2015).

44. Zhang, Y. et al. Neuronal TORC1 modulates longevity via AMPK and cell nonautonomous regulation of mitochondrial dynamics in C. elegans. eLife 8, e49158 (2019).

45. Lee, S.-J. & Kenyon, C. Regulation of the Longevity Response to Temperature by Thermosensory Neurons in Caenorhabditis elegans. Curr. Biol. 19, 715–722 (2009).

46. Leiser, S. F., Begun, A. & Kaeberlein, M. HIF-1 modulates longevity and healthspan in a temperature-dependent manner. Aging Cell 10, 318–326 (2011).

47. Sado, M. et al. Protective effect against Parkinson’s disease-related insults through the activation of XBP1. Brain Res. 1257, 16–24 (2009).

48. Casas-Tinto, S. et al. The ER stress factor XBP1s prevents amyloid-β neurotoxicity. Hum. Mol. Genet. 20, 2144–2160 (2011).

49. Vidal, R. L. et al. Targeting the UPR transcription factor XBP1 protects against Huntington’s disease through the regulation of FoxO1 and autophagy. Hum. Mol. Genet. 21, 2245–2262 (2012).

50. Brenner, S. The genetics of Caenorhabditis elegans. Genetics 77, 71–94 (1974).

51. Timmons, L., Court, D. L. & Fire, A. Ingestion of bacterially expressed dsRNAs can produce specific and potent genetic interference in Caenorhabditis elegans. Gene 263, 103–12 (2001).

52. Vicencio, J., Martínez-Fernández, C., Serrat, X. & Cerón, J. Efficient Generation of Endogenous Fluorescent Reporters by Nested CRISPR in Caenorhabditis elegans. Genetics 211, 1143–1154 (2019).

53. Paix, A. et al. Scalable and versatile genome editing using linear DNAs with microhomology to Cas9 Sites in Caenorhabditis elegans. Genetics 198, 1347–56 (2014).

