## Supplemental material for "Olfactory chemosensation extends lifespan through TGF-β signaling and UPR activation"

### Supplemental Information

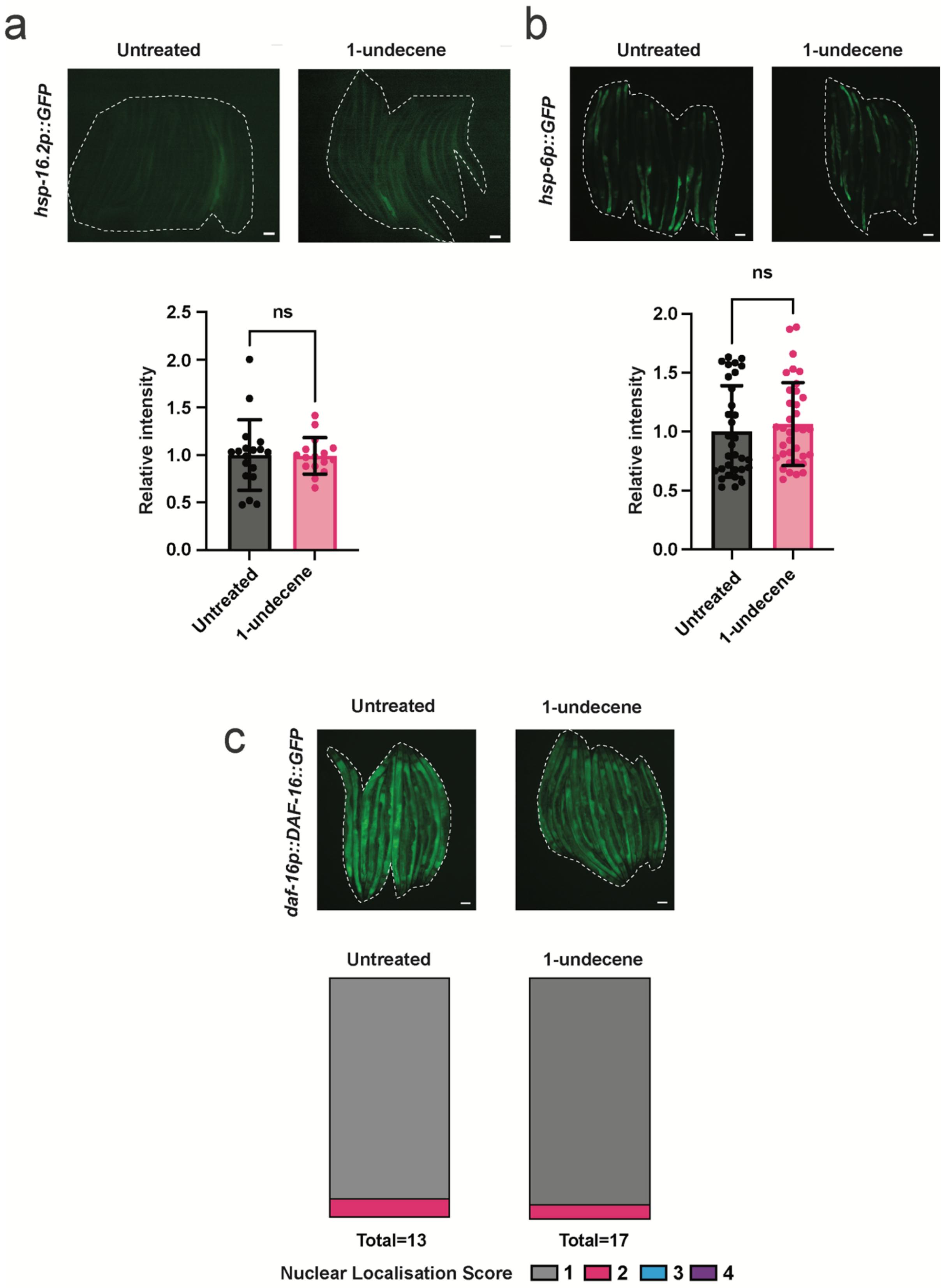

**Extended Data Fig. S1: 1-undecene exposure does not activate other stress response pathways. a,b,** Representative fluorescence microscopy images and quantification of **a,** *hsp-16.2p::GFP* and **b,** *hsp-6p::GFP* fluorescence. These experiments were repeated three times with at least 10 worms per group. Ns, not significant (unpaired Student's t test). Scalebars, 200 µm. **(C)** Representative fluorescence microscopy images and quantification of the subcellular localization of DAF-16::GFP in worms expressing a *daf-16p::DAF-16::GFP* transgene. Worms were scored based on the number of intestinal cells that presented nuclear GFP localization, 1 = 0 (cytosolic GFP only), 2 = 2-4 cells, 3 = 5-8 cells, 4 = more than 8 cells. This experiment was repeated three times with more than 10 worms per group. Scalebars, 200 µm.

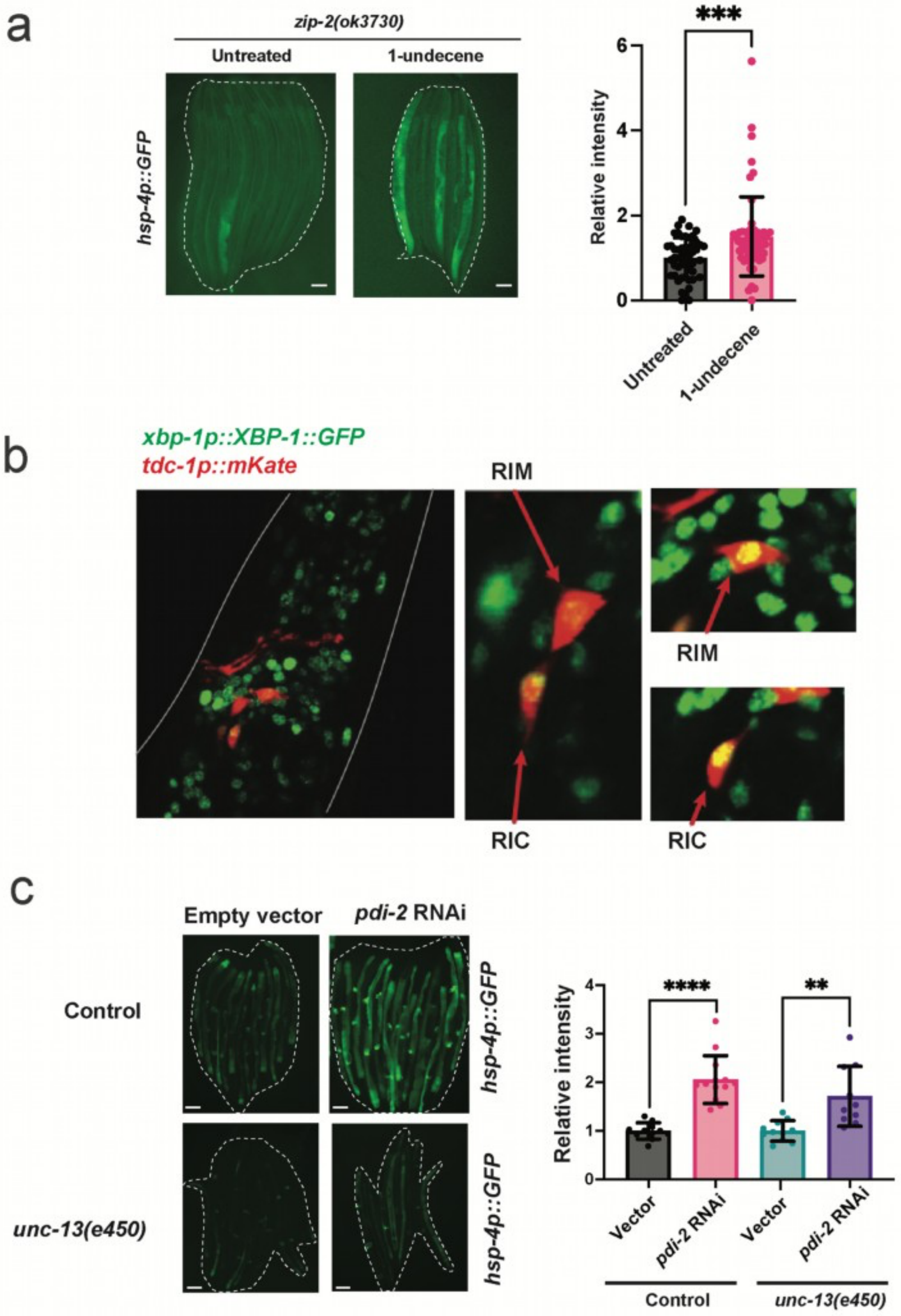
a
zip-2(ok3730)
Untreated
1-undecene
hsp-4p::GFP
***
Relative intensity
6
4
2
0
Untreated
1-undecene
b
xbp-1p::XBP-1::GFP
tdc-1p::mKate
RIM
RIM
RIC
RIC
c
Empty vector
pdi-2 RNAi
Control
unc-13(e450)
hsp-4p::GFP
hsp-4p::GFP
****
**
Relative intensity
4
3
2
1
0
Vector
pdi-2 RNAi
Vector
pdi-2 RNAi
Control
unc-13(e450)

**Extended Data Fig. S2: Activation of the UPR$^{ER}$ in by 1-undecene odor does not require the immune response regulator ZIP-2 and occurs in RIM/RIC interneurons. a**, Representative fluorescence microscopy images and quantification of *hsp-4p::GFP* fluorescence in *zip-2(ok3730)* animals with or without exposure to 1-undecene odor for 12 hours. ***$P$<0.001 (unpaired Student's t test). This experiment was repeated three times with at least 10 worms per group. Scalebars, 200 µm. **b,** Representative image of worms expressing *tdc-1p::mKate; xbp-1p::xbp-1::GFP* transgenes exposed to 8 hours of 1-undecene odor. Scalebars, 10 µm. **c,** Representative fluorescence microscopy images and quantification of *hsp-4p::GFP* fluorescence in *hsp-4p::GFP* animals with or without an *unc-13(e450)* mutant background grown from L1 larval stage on NGM plates containing bacteria harboring empty vector (L4440) or *pdi-2* RNAi. Data were normalized by samples treated with Vector only. The experiment was repeated twice with at least 8 animals per group. **$P$<0.01, ****$P$<0.0001 (unpaired Student's t test). Scalebars, 200 µm.

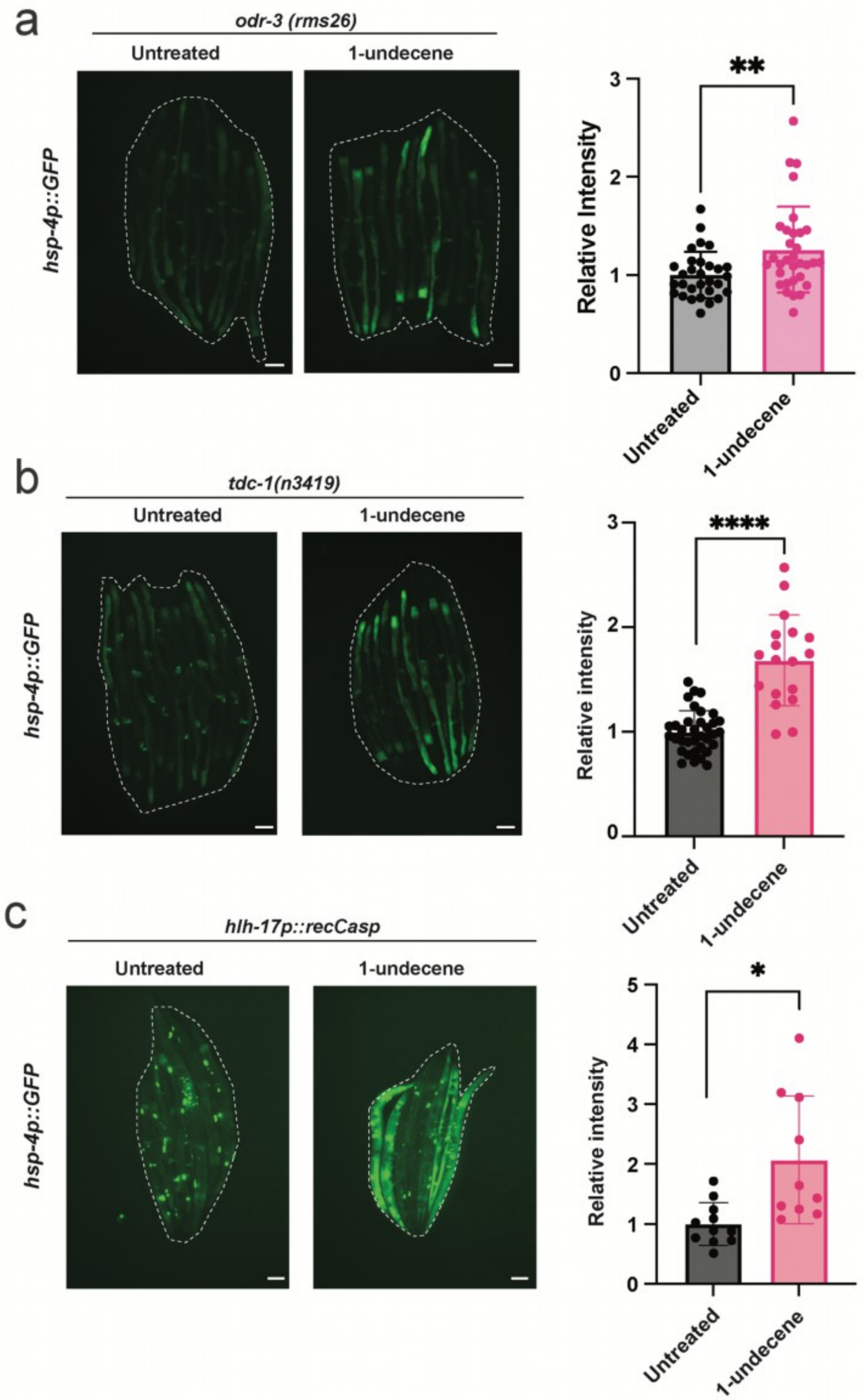
a
odr-3 (rms26)
Untreated
1-undecene
hsp-4p::GFP
**
Relative Intensity
3
2
1
0
Untreated
1-undecene
b
tdc-1(n3419)
Untreated
1-undecene
hsp-4p::GFP
****
Relative intensity
3
2
1
0
Untreated
1-undecene
c
hlh-17p::recCasp
Untreated
1-undecene
hsp-4p::GFP
*
Relative intensity
5
4
3
2
1
0
Untreated
1-undecene

**Extended Data Fig. S3: ODR-3, tyramine synthesis, and CEPsh glia are not required for $UPR^{ER}$ activation by 1-undecene odor. a,b,c,** Representative fluorescence microscopy images and quantification of *hsp-4p::GFP* fluorescence in animals with **a,** *odr-3(rms31)*, **b,** *tdc-1(n3419)*, and **c,** *nsIs180[hlh-17p::recCaspase-3, unc-122p::GFP]* backgrounds with or without exposure to 1-undecene for 12 hours. Experiments were repeated three times with at least 8 worms per group. **P*<0.05, ***P*<0.01, *****P*<0.0001 (unpaired Student's t test). Scalebars, 200 µm.

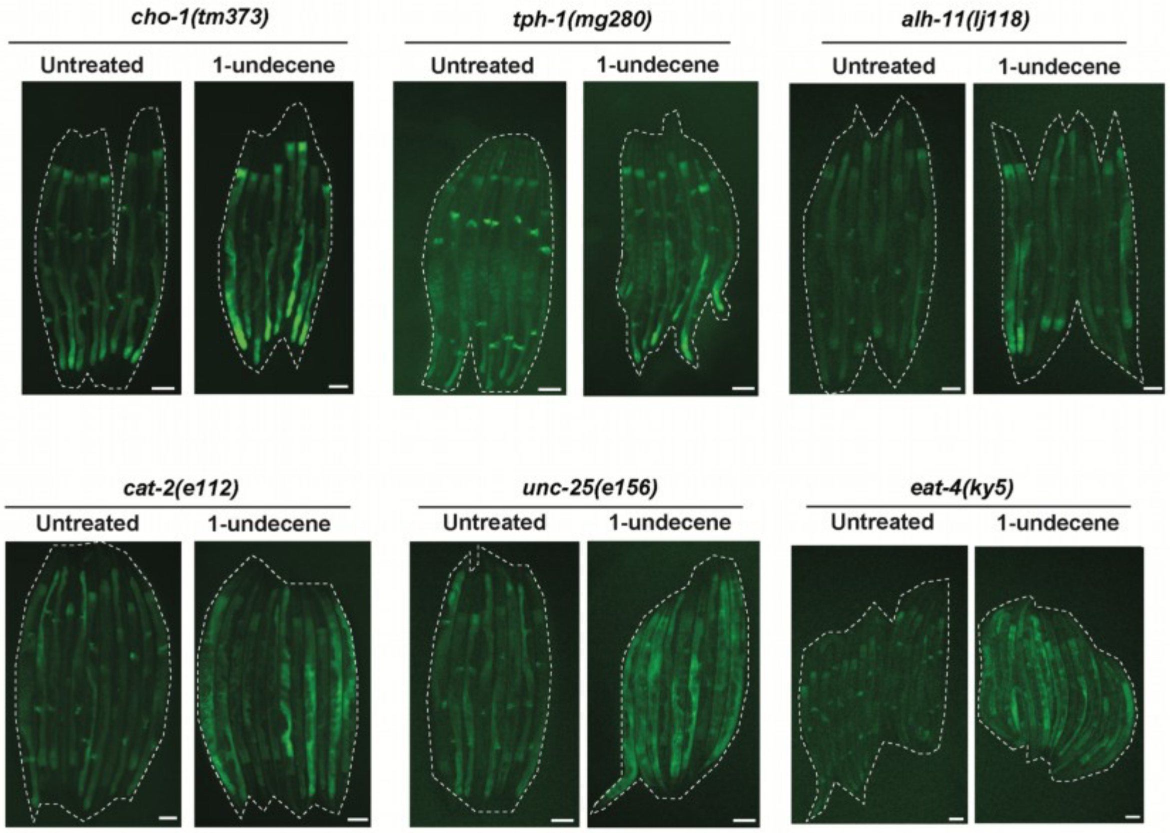
cho-1(tm373)
Untreated
1-undecene
tph-1(mg280)
Untreated
1-undecene
alh-11(lj118)
Untreated
1-undecene
cat-2(e112)
Untreated
1-undecene
unc-25(e156)
Untreated
1-undecene
eat-4(ky5)
Untreated
1-undecene

**Extended Data Fig. S4: A range of neurotransmitters are dispensable for activation of the *hsp-4p::GFP* reporter transgene by 1-undecene exposure.** Representative fluorescence microscopy images of *hsp-4p::GFP* fluorescence in *cho-1(tm373)*, *tph-1 (mg280), alh-11 (lj118), cat-2*(*e1112*), *unc-25(e156)*, and *eat-4(ky5)* backgrounds after exposure or no exposure to 1-undecene for 12 hours. Experiments were repeated three times with at least 8 worms per group. Scalebars, 200 µm.

**Supplementary Table S1. Survival information for lifespan experiments.**

| Strain | Treatment | Median lifespan (days) | Control | Median lifespan (days) | *P* value (Mantel-Cox) |
|---|---|---|---|---|---|
| N2 | + 1-undecene | 20 | - 1-undecene | 18 | 0.0227 |
| N2 | + 1-undecene | 20 | - 1-undecene | 18 | 0.0357 |
| N2 | + 1-undecene | 26 | - 1-undecene | 21 | 0.0185 |
| *xbp-1(zc12)* | + 1-undecene | 14 | - 1-undecene | 14 | 0.7015 |
| *xbp-1(zc12)* | + 1-undecene | 18 | - 1-undecene | 20 | <0.0001 |
| *xbp-1(zc12)* | + 1-undecene | 22 | - 1-undecene | 22 | 0.0066 |

**Supplementary Table S2. List of *C. elegans* strains used.**

| ***Caenorhabditis elegans* strains** | | |
|---|---|---|
| Wild type | CGC | N2 |
| *zcIs4 [hsp-4::GFP]* V | CGC | SJ4005 |
| *ire-1(zc14)* II*; zcIs4 [hsp-4::GFP]* V | CGC | SJ30 |
| *zIs356 [daf-16p::daf-16a/b::GFP + rol-6(su1006)]* IV | CGC | TJ356 |
| *rmIs132 [unc-54p::Q35::YFP]* | CGC | AM140 |
| *dvIn70* [*hsp-16-2p::GFP*; *rol-6*] | CGC | CL2070 |
| *daf-11(m47)* V | CGC | DR47 |
| *rmIs110 [F25B3.3p::Q40::YFP]* | Özbey *et al.,* 2019 | AGD1397 |
| *uthIs393 [vha-6p::Q40::YFP+rol-6(su1006)]* | Özbey *et al.,* 2019 | AGD1395 |
| *rmsIs9* [*daf-7p::xbp-1s::unc-54* 3'UTR, *myo-3p::mKate*]; *zcIs4 [hsp-4p::GFP]* V | Özbey *et al.,* 2020 | RCT206 |
| *rmsIs8 [xbp-1p::xbp-1::GFP]* | Özbey *et al.,* 2020 | RCT21 |
| *rmsIs7* [*tdc-1p::mKate2::let-858* 3'UTR, *cc::GFP*]; *rmsIs8* [*xbp-1p::xbp-1::GFP*] | Özbey *et al.,* 2020 | RCT192 |
| *xbp-1(zc12)* III | Taylor & Dillin, 2013 | AGD1049 |
| *xbp-1(zc12)* III; *zcIs4 [hsp-4::GFP]* V | Taylor & Dillin, 2013 | AGD972 |
| *unc-13 (e450)* I; *zcIs4 [hsp-4::GFP]* V | Taylor & Dillin, 2013 | AGD1137 |
| drcSI7 [*daf-7p::Venus*] | Tullet Lab, U. of Kent | JMT50 |
| *zip-2(ok3730)* III*; zcIs4 [hsp-4::GFP]* V | This study | RCT369 |
| *unc-31 (e928)* IV*; zcIs4 [hsp-4::GFP]* V | This study | RCT370 |
| *tdc-1 (n3419)* II*; zcIs4 [hsp-4::GFP]* V | This study | RCT66 |
| *tph-1 (mg280)* II; *zcIs4 [hsp-4::GFP]* V | This study | RCT371 |
| *cat-2*(*e1112*) II; *zcIs4 [hsp-4::GFP]* V | This study | RCT372 |
| *unc-25(e156)* III; *zcIs4 [hsp-4::GFP]* V | This study | RCT373 |
| *eat-4(ky5)* III; *zcIs4 [hsp-4::GFP]* V | This study | RCT374 |
| *nsIs180 [hlh-17p::recCaspase-3, unc-122p::GFP]; zcIs4 [hsp-4::GFP]* V | This study | RCT375 |
| *cho-1(tm373)* IV; *zcIs4 [hsp-4::GFP]* V | This study | RCT189 |
| *alh-11 (lj118)* III; *zcIs4 [hsp-4::GFP]* V | This study | RCT376 |
| *daf-1 (m40)* IV; *zcIs4 [hsp-4::GFP]* V | This study | RCT67 |
| *daf-7 (e1372)* III; *zcIs4 [hsp-4::GFP]* V | This study | RCT68 |
| *odr-3(rms31); zcIs4 [hsp-4::GFP]* V | This study | RCT377 |

**Supplementary Table S3. List of qPCR primers used in this work.**

| | SOURCE | IDENTIFIER |
|---|---|---|
| **qPCR Primers** | | |
| GTTCCCGTGTTCATCACTCAT | Sigma | *pmp-3* F |
| ACACCGTCGAGAAGCTGTAGA | Sigma | *pmp-3* R |
| CTGCTGGACAGGAAGATTACG | Sigma | *cdc-42* F |
| CTCGGACATTCTCGAATGAAG | Sigma | *cdc-42* R |
| GTCGCTTCAAATCAGTTCAGC | Sigma | *Y45F10D.4* F |
| GTTCTTGTCAAGTGATCCGACA | Sigma | *Y45F10D.4* R |
| CGTGCCTTTGAATCAGCAGTG | Sigma | *xbp-1*_s F |
| CGAGGTGTCCATCTTCTTGTT | Sigma | *xbp-1*_s R |
| CAGATGAAAACTCAAATCGCC | Sigma | *hsp-4* F |
| GGTTGCTTCCGAGCCACTCAA | Sigma | *hsp-4* R |
| CTTTATTCGCTCCCTAACCGT | Sigma | *Y41C4A.11* F |
| CTGCTCCATTCTCCACTCGTT | Sigma | *Y41C4A.11* R |
| TTCTACACCGCCAAAGATCC | Sigma | *daf-7* F |
| CTGTGAGTGTGGCCTGAAGA | Sigma | *daf-7* R |
| AGCCGGCGATGTAAGCTAAG | Sigma | *odr-3* F |
| ACCCTTAAGCGCACTCTCAAC | Sigma | *odr-3* R |

**Supplementary Table S4. List of CRISPR primers and oligonucleotides used in this work.**

| | SOURCE | IDENTIFIER |
|---|---|---|
| **CRISPR Oligos** | | |
| GCTACCATAGGCACCACGAG | IDT | *dpy-10* cRNA |
| AAAATTCGGAAGGTAACGCG; C TGCATTTACCGTTGGAAAA | IDT | *odr-3* cRNAs |
| CACTTGAACTTCAATACGGCAAGATGAG AATGACTGGAAACCGTACCGCATGCGG TGCCTATGGTAGCGGAGCTTCACATGGC TTCAGACCAACAGCCTAT | IDT | *dpy-10* Repair Template |
| caattactcatagatcattgtttttttagATATGGGCTCA TGCCAGAGCAATGAAAATTCGGAAGGTA ACAAATGGACAAAAAGAAGCAGAAAAGG CAATAGTTATGAAAGTACAGGAAAATGGAGAAG AAGGAGAAGCACTGACAGAAGAAGTTTCGAAA GCAATTCAATCGTTATGGGCAGATCCTGGCGTGAAGAA | IDT | *odr-3* Repair Template |
